# Vector analysis of steerable mechanical tension across nuclear lamina

**DOI:** 10.1101/462275

**Authors:** TingTing Chen, HuiWen Wu, YuXuan Wang, JinJun Shan, JiaRui Zhang, HuanHuan Zhao, Jun Guo

## Abstract

The nucleus is the most prominent organelle in eukaryotic cells, and its deformation depends on interactions between the nuclear lamina (NL) and cytoskeleton structural tensions. The structural tensions can be quantified at a pico-Newton (pN) level using a genetically encoded optical probe. In living cells, NL tensions countered the 4.26pN resting strain imposed competitively by cytoskeletal tension. The depolymerization of microfilaments or microtubules drove an aberrant increase in outward osmotic pressure through the production of mass protein-nanoparticles. The osmotic pressure also served as a directional converter of inward cytoskeletal force, and contributed to the outward expansion of NL via the passive pull of intermediate filaments (IFs). The NL, but not IFs, can remotely detect extracellular osmosis pressure alterations, which are closely associated with highly polarized microfilament and microtubule structures and their directional force activities. The oxidative-induced increase of NL tension results from intracellular hyper-osmosis, associated closely with protein-nanoparticles production elicited by cofilin and stathmin activation. These data reveal that intracellular steerable forces interact direction-dependently to control NL tension in terms of their magnitude and vectors.

## INTRODUCTION

The nuclear lamina (NL) is a thin filamentous meshwork that provides stabilizing mechanical support to the nucleus and hinges on the appropriate feedback of mechanical properties^1-4^. As the central nuclear skeletal hub and coordinator, the NL can be subjected to significant forces generated by the cytoskeleton and can adjust the shape of the nucleus during the development, differentiation, invasion, apoptosis and senescence of cells ^5-8^.

Nuclear deformation mainly proceeds through a balance of cytoskeletal forces generated by MFs and MTs that pull, push and shear the nucleus, and IFs that may passively resist nuclear decentering and deformation^6^. The LINC (linker of nucleoskeleton and cytoskeleton) complex facilitates internal cell interconnectivity between MFs, MTs, and IFs to the nucleus, involving in the nucleus to directly perceive and integrate transmission of cytoskeletal structural tension^9-11^. MFs, MTs, and IFs appear to act as “tensile guidewires” that anchor the nucleus in place and coordinate changes in nucleus deformation^10, 12^. However, understanding the contributions of these different structural forces within complex nuclear mechanics is a very challenging task.

Internal load-bearing cytoskeletal structural tension can also be regulated by external osmotic pressure (OP). OP can be generated from concentration differences among ion and colloid particles between the cytoplasm and extracellular fluid^13^. Osmotic pressure can induce outward or inward tension on the nucleus through the pulling or pushing of cytoskeletal IFs^14, 15^. IF tension exerts pulling forces on the nucleus, which serves as a “piston” with the function of osmotic pressure. Furthermore, osmotic pressure-induced IF tension can be reversed by MF and MT tensions in the phenomenon of regulatory volume decrease (RVD) ^14^.

MFs and MTs, unlike IFs, are highly polarized^16^. The organization of the MF and MT network in cells, where plus ends are found adjacent to the membrane and minus ends are located toward the cytoplasm, imply their role in pulling or pushing plasma membrane and nuclear envelope^17-19^. Myosin has a role in prograde transport along MFs toward the membrane. Most kinesin proteins move towards the microtubule plus ends^15, 20,^ whereas cytoplasmic dynein locates along the minus-end-directed motor in the cell^21, 22^. The direction of cytoskeletal structural tensions is dependent upon the organization of the cytoskeletal meshwork and the direction of the molecular motor^16, 23^. Meanwhile, tension due to MFs and MTs could be eliminated if MFs and MTs are depolymerized into actin and tubulin monomers or macromolecular polymers of size 1–100 nm^24-26^. Generation of these nanoparticle was thought to result in the colloid OP. However, the mechanisms that underpin the interactions between external forces and internal stresses in nucleoskeletal alterations remains poorly understood.

Nuclear deformation can be induced in live cells in response to chemical trigger signals, which may be associated with senescence^7,^ oxidative stress^27, 28,^ and apoptosis-related changes^29, 30^. These activities have all been identified as mechanosensitive elements that can activate cytoskeletal structural tension^31, 32^. Tension can be rapidly redistributed at a subcellular level and can reprogram cell shape to maintain balanced cell integrity^33-36^. However, in the absence of a method to measure the forces across structural proteins at a subcellular level, it has been difficult to define how nuclear-laminal tension influences nuclear morphological and functional changes. Consequently, subcellular tensions, including their magnitude and direction, remain poorly understood in response to chemical stimuli.

With this in mind, we recently developed genetically encoded FRET (fluorescence resonance energy transfer)-based tension probes. We clarify and extend these previously reported findings to provide mechanistic insights into tension interactions in living cells and how these tensions influence laminar function through change of their magnitude and orientation under oxidative stress.

## RESULTS

### 1. Measurements of FRET in intracellular structural tension probes and quantity of the tension in nuclear lamina

It is well known that intracellular tension is dependent upon cytoskeletal transmission. To investigate intracellular structural tensions and their interactions, the dipole-orientation-based FRET probe can be incorporated into cytoskeletal proteins according to previous report ^37-39^, like Lamin B1–cpstFRET–Lamin B1 (LBcpLB), actin–cpstFRET–actin (AcpA) and α-tubulin–cpstFRET–β-tubulin (TcpT), and vimentin–cpstFRET–vimentin (VcpV), and provide real-time tension measurements of NL, MFs, MTs, and IFs in live cells (Fig. 1A). CpstFRET is primarily modulated by the angle between the donor and the acceptor, which is parallel at rest and is expected to twist toward to a perpendicular configuration under tension. Colocalization experiments with tension probes (FRET fluorescence) showed high spatial correlation between these probes (yellow) and corresponding cytoskeletons (red), accompanied with a calculated Pearson’s correlation coefficient (PCC) of > 0.86 (Fig. 1B), suggesting that the expressed probes could be incorporated into cytoskeleton filaments and bridge adjacent filaments. Western blot analysis was used to define the molecular weight of each probe (sFig. 2A). Meanwhile, no significant differences in terms of the cell-cycle, apoptosis rate, or cytoskeletal structure were observed in cells transfected with these tension probes compared to controls (sTab. 1, sFig. 1, and sFig. 2B). Thus, insertion of cpstFRET into the cytoskeleton did not significantly affect cell physiological processes.

**Figure 1.**
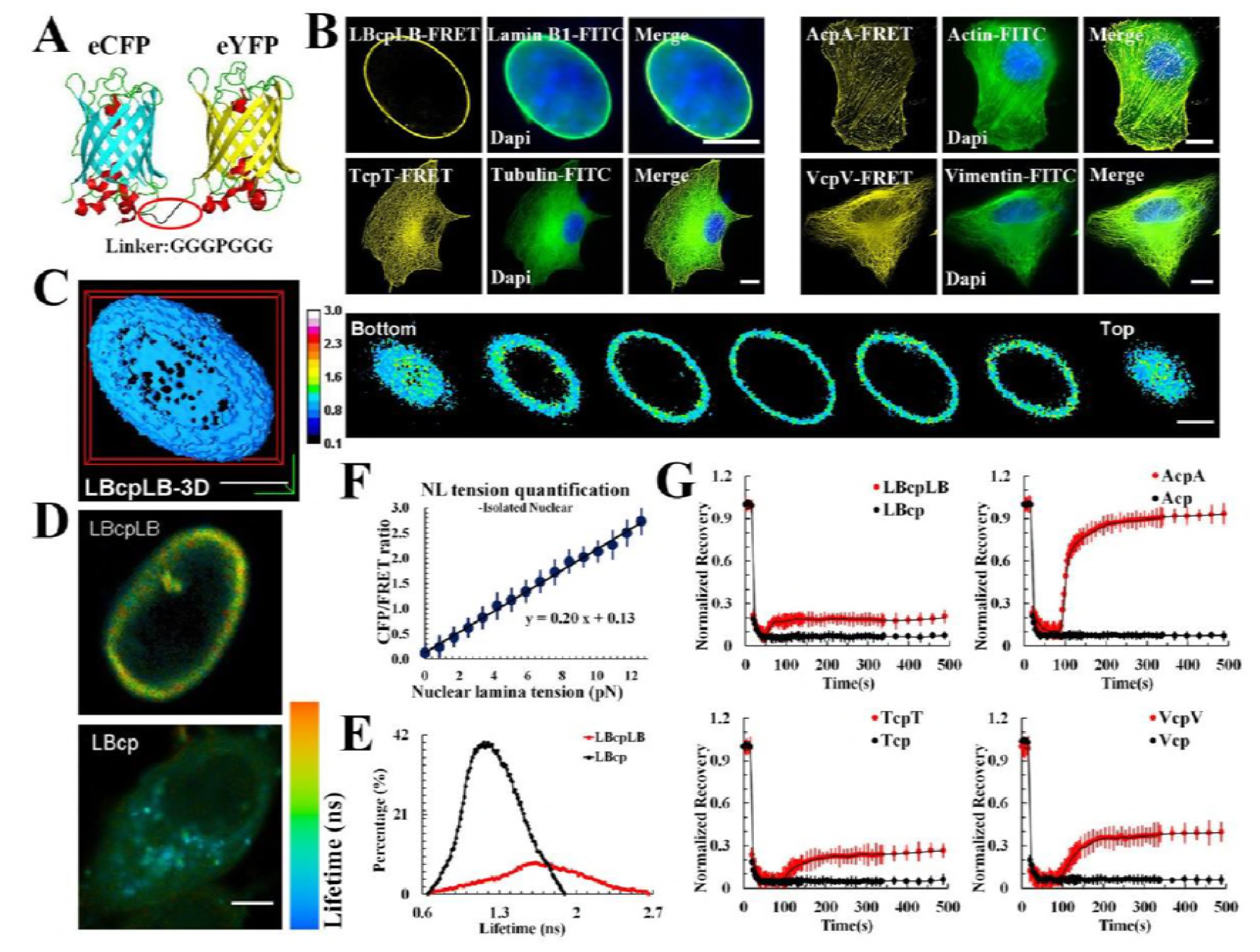
LBcpLB, AcpA, TcpT, and VcpV tension probe (cpst FRET) construct. (A) The FRET-based probe module (cpstFRET) was comprised of donor (cpVenus) and acceptor (cpCerulean) separated by an amino acid sequence (3aa–Pro–3aa). The cpVenus lies parallel to the cpCerulean at rest. When external force across are applied, the cpstFRET generates a certain angle, reducing the Förster resonance energy transfer (FRET) efficiency (f: external force)^47^. Lamin B1 at the C-terminal of cpstFRET was cloned, creating pEG–lamin B1–cpstFRET (LBcp). cpstFRET was inserted into two the lamin B1 gene to create pEG–lamin B1–cpstFRET–lamin B1 (LBcpLB). Similarly, AcpA, TcpT and VcpV were constructed. (B) Confocal photomicrographs of MCF-7 cells expressing LBcpLB, AcpA, TcpT, and VcpV tension probes, respectively. FRET fluorescence (yellow) overlaid with images of FITC-stained cytoskeleton fluorescence (green). Colocalization of tension probes and FITC fluorescence appears as merged images. (C) Three-dimensional (3D) constructions and representative 3D images of LBcpLB tension (n = 9). (D) Fluorescence lifetime images of MCF-7 cells expressing LBcpLB or LBcp. (E) Fluorescence lifetime histograms from LBcpLB nuclear lamina (n = 10) and LBcp (n = 10) expressing MCF-7 cells. (F) The linear relationship between osmotic pressure and CFP/FRET ratio in *in vitro* nuclei: Y = 0.20X + 0.13 (n = 11). (G) Normalized average fluorescence recovery of LBcpLB, AcpA, TcpT, VcpV and their respective controls versus time (n = 10). The calibration bar was set from 0.1 to 3.0. Scale bar, 10 μm. All error bars represent the SEM.

To verify the accuracy of the fluorescence and efficiency of the FRET in the tension probe, we transfected MCF-7 cells with the probes mentioned above and examined them via laser confocal scanning microscopy. Fluorescence lifetime imaging microscopy (FLIM) technique (Fig. 1D and 1E), acceptor bleaching (AB) analysis (sFig. 4A), fluorescence recovery after photobleaching (FRAP) measurements (Fig. 1G), and three-dimensional cell reconstructions (Fig. 1C) were employed, and the data showed that LBcpLB, AcpA, TcpT and VcpV are effective probes in terms of authenticity, efficiency, and intracellular dynamics. Compared with their respective controls, LBcp, Acp, Tcp and Vcp, the FRET indices (eCFP/FRET) were very high compared with the tension probes (sFig. 3).

To quantify the tension in the nuclear lamina, we used isolated MCF-7 nuclei ^40^ with the LBcpLB probe. The isolated nucleus reacted to the hypotonic pressure could maintain its rounded morphology (600→300 mOsm/kg, sFig. 4B; 300→0 mOsm/kg, sFig. 4D). The eCFP/eYFP fluorophore ratio changed rapidly under different osmotic pressure cycles (sFig. 4C and 4E). These measurements were used to estimate the force of cpstFRET and to calculate forces across LBcpLB in living cells using osmotic pressure data (sFig. 4F). The conversion relationship between the osmotic pressure values and the induced NL tension are shown in sFig. 5. Combined, these results indicate that NL tension was linearly correlated to the eCFP/eYFP ratio (Fig. 1F) and that the LBcpLB probe was most sensitive between 0.0 to 12.6 pN. The average force for a stationary LBcpLB probe within a living cell was measured to be ~4.3 pN.

### 2.

Depolymerisation of MFs and MTs is involved in up-regulation of nuclear lamina tension through the pulling force of IFs and the protein-nanoparticle induced osmotic pressure.

The generation and transduction of intracellular mechanical tension relies on intracellular cytoskeleton architecture ^41, 42^. To investigate the changes in intracellular mechanical activities induced by cytoskeletal structure alterations, cytochalasin D (Cyto D) and nocodazole (Noc) were used to depolymerize MFs and MTs, respectively. Filiform MFs and MTs structure turned into punctiform granules in response to depolymerizing agents (Fig. 3A). NL and a cytoskeleton (MF, MT or IF) tension probe were cotransfected into MCF7 cells. Neither depolymerizing agents influenced cytoskeletal protein content (sFig. 2C). Figure 2A and 2B shows MF or MT depolymerization resulted in temporal increment of IF-and NL-tension (15 min), accompany with upregulation of MT or MF tension. Among these, the NL tension was higher than either of the cytoskeletal structural tensions.

**Figure 2.**
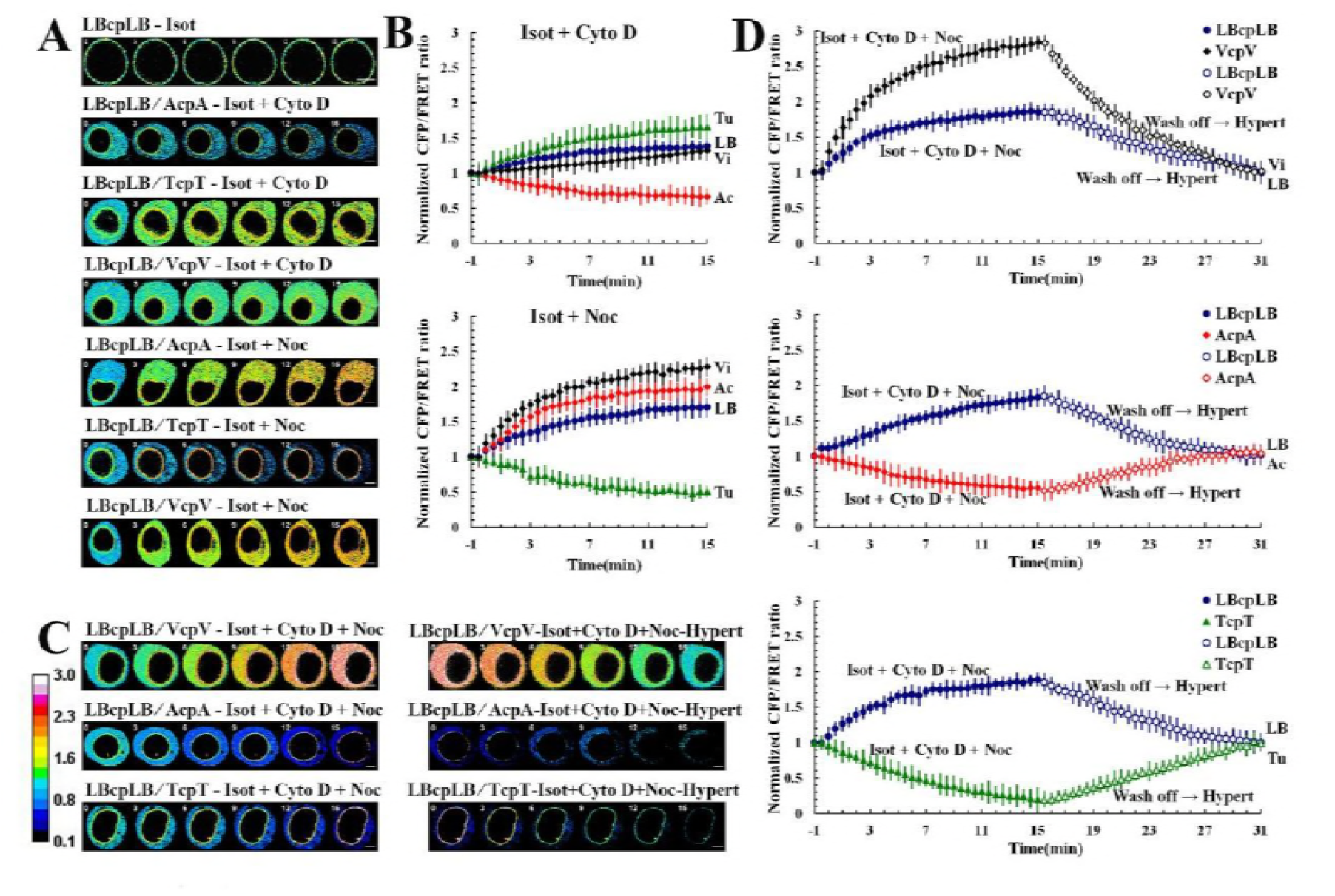
Effect of cytoskeletal depolymerization agents on cytoskeletal structural tension and nuclear lamina tension under isotonic conditions and the response to hyper-osmotic pressure. The cpstFRET probe module was used to measure changes in cytoskeletal and nuclear lamina tensions in MCF-7 cells. The graph plots are normalized to CFP/FRET signals for untreated cells and those treated with MF (Cyto D) and MT (Noc) depolymerization agents under isotonic conditions or in recuperative responses to hyper-osmotic pressure (n = 9). (A) Under isotonic conditions, MCF-7 cells were transfected with LBcpLB alone (row 1) and were also transfected with LBcpLB together with AcpA, TcpT, and VcpV individually (row 2 –7). NL, MF, MT, and IF tensions in MCF-7 cells were analyzed after exposure to cyto D (row 2–4) and Noc (row 5-7). (B) Normalized CFP/FRET signals corresponding to NL (blue), MF (red), MT (green), and IF (black) tensions versus time in response to Cyto D or Noc stimuli, under isotonic conditions versus (A). (C) Under isotonic conditions, MCF-7 cell NL, MF, MT, and IF tensions were analyzed after exposure to Cyto D and Noc simultaneously. Then, MCF-7 cells were exposed to hyper-osmotic conditions by washing off the bath from isotonic (300 Osm/kg) to hypertonic (540 Osm/kg). (D) Normalized CFP/FRET signals corresponding to NL, MF, MT, and IF tensions versus time in response to the two agents simultaneously, under isotonic (solid points) and hypertonic conditions (hollow points). The calibration bar was set from 0.1 to 3.0. Scale bar, 10 μm. All error bars represent the SEM.

However, when MFs and MTs were simultaneously depolymerized, IF- and NL-dependent tension collectively heightened, reaching the same level after 15 min (Fig. 2C, left). Then, we challenged the cells with hyper-osmotic conditions by washing off the bath from isotonic saline conditions (300 Osm/kg) to hypertonic (540 Osm/kg) conditions. The cell bodies and nuclei reverting to their original volume confirmed the hypertonic environment inside the cells. IF-dependent tension reverted to the initial values and remained equal to the NL tension (Fig. 2C, right). MF- and MT-dependent tensions partly recovered due to depolymerization (Fig. 2C, 2D, and Movie S1-3). We also observed that both the cell and nuclear volume enlarged distinctly. A similar phenomenon was also observed when MF and MT were depolymerized separately or jointly (Fig. 3C and sFig. 7A). Taken together, these data suggest that the increment in NL and IF tension were caused by outward swelling tension, likely exerted by actin and tubulin particles from MF and MT depolymerization.

**Figure 3.**
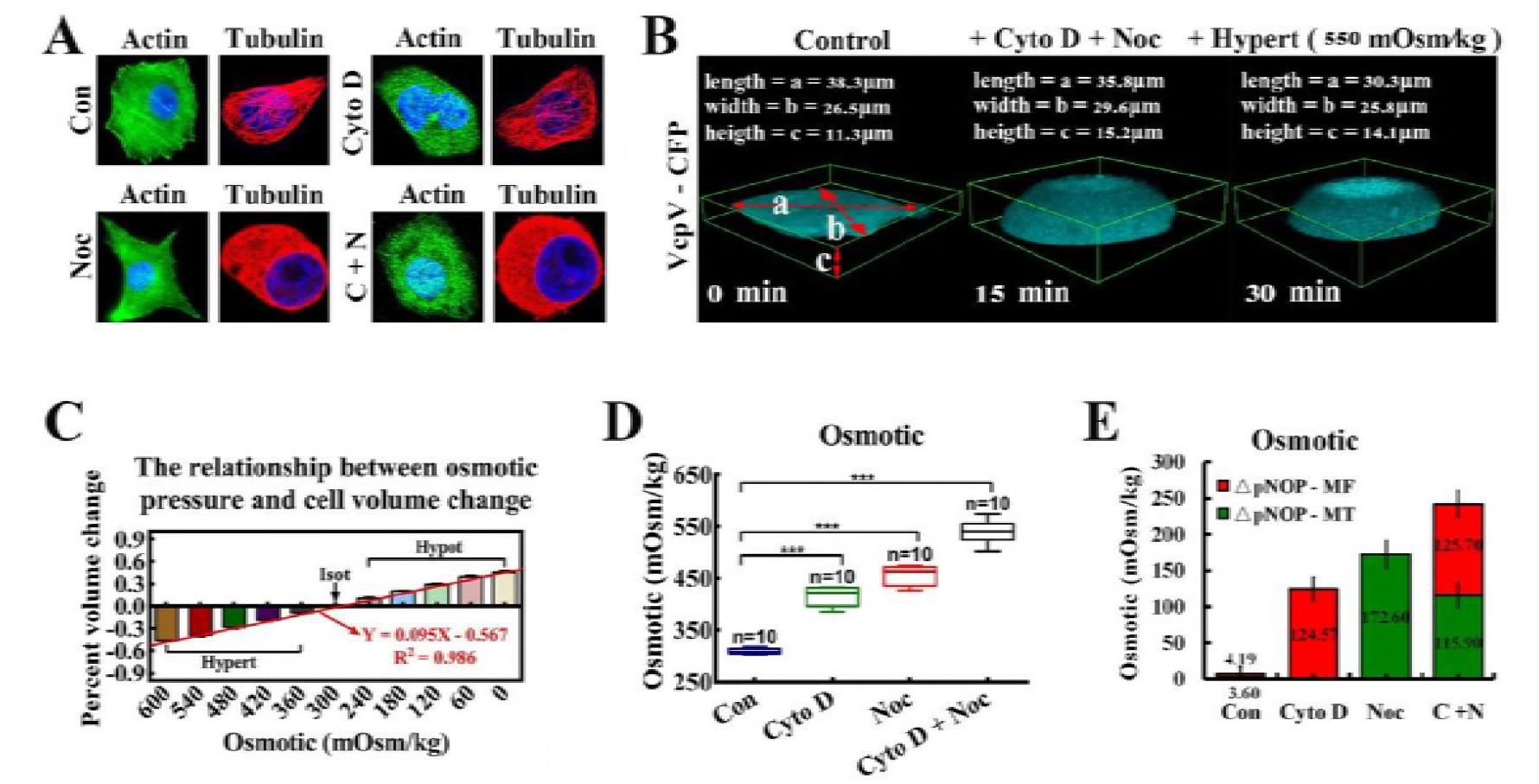
Intracellular osmotic pressure composition changes in response to MF and MT depolymerization agents. (A) MCF-7 cells were treated with Cyto D and Noc, respectively and simultaneously. β-actin and α-tubulin were stained respectively. The cells were also stained with DAPI to localize the nuclei. The images were captured through confocal laser microscopy after immunofluorescence staining (n = 10). (B) Cell deformation was induced by the osmotic pressure difference across the membrane. The extracellular osmotic pressure and the percent of cell volume change were measured and their linear relationship was determined: Y = 0.095X – 0.567 (n = 10), R^2^ = 0.986. (C) Three-dimensional (3D) reconstructions of MCF-7 cells (n = 9). The cells volume was calculated by using the formulae: V = 4π/3*(a/2)*(b/2)*(c/2). (D) Cells were treated with Cyto D and Noc, respectively and simultaneously. The linear relationship was applied to calculate the intracellular osmotic pressure (***: p < 0.001, ns: p > 0.05, Tukey-b test, n = 10). (E) This linear relationship was applied to calculate the proportional changes in intracellular protein-nanoparticle induced pressure (ΔpNOP) in response to different depolymerizing agents. Count rates were measured after treatment with depolymerizing agents (n = 10). MF or MT depolymerization (individually or simultaneously) can cause ΔpNOP gain. Scale bar, 10 μm. All error bars represent SEM.

More efforts were made to confirm this hypothesis. Direct measurements of intracellular osmotic pressure were made by osmometry. Compared to control cells, when depolymerizing agents were applied to the test cells, significant increases in intracellular osmotic pressure were recorded (Fig. 2E, 3C and 3D). Meanwhile, nanometre scale experiments confirmed the presence of actin (5–6 nm) and tubulin (7–8 nm) nanoparticles (sFig. 7B). Additionally, western blot analysis revealed that the depolymerizing agents did not alter the total protein content in living cells (sFig. 7C). Meanwhile, reductions in MF and MT tensions through calcium elimination or ATP depletion did not elicit changes in IF tension (sFig. 6). Together, these outcomes verified the generation of intracellular colloid osmotic pressure through MF and MT depolymerisation and production of mass protein nanoparticles, which led to the enhancement of IF and NL tension. Moreover, intracellular nanoparticle-induced osmotic pressure was quantified. The linear relationship between the osmotic pressure and percent cell volume change is illustrated in Fig. 3B: Y = 0.095X – 0.567, R^2^=0.986 (the osmotic pressure gradient significantly caused the changes of cellular morphology, 600→0 mOsm/kg). Treatment with Cyto D, Noc or both appreciably increased intracellular protein-nanoparticle-induced osmotic pressure (Fig. 3D and 3E). This result suggests that protein-nanoparticle induced osmotic pressure is an essential contributor to overall osmotic pressure in living cells and increases though the large-scale production of β-actin (5–6nm) and α/β-tubulin (7–8nm) nanoparticles (1–10% of the total cell protein) due to cytoskeleton depolymerization. Therefore, the intracellular osmotic pressure was dependent upon nanoarticle-induced osmotic pressure and ionic osmotic pressure, which maintained the dynamic osmotic equilibrium under normal physiological conditions.

### 3. Vector analysis of cytoskeletal structural forces involving in nuclear lamina tension in response to external osmotic-pressure stimuli

To explore the role of osmotic pressure in cytoskeleton structural tensions, extracellular hypotonic or hyper-osmotic stimuli were employed, and magnitude and direction of the structural tensions were analysed. Under hypotonic conditions, MT-, MF-, IF-, and NL-tensions increased as illustrated in Fig. 4A and 4B, and IF tension was much lower than NL tension. To determine if MF- and MT- tension influence NL- and IF-tension, we employed BMD, Cilio D, and SB-715992 to inhibit myosin, dynein, and kinesin motion, respectively (Fig. 4D, 4F, 4K and 4M). The three molecular motor inhibitors did not affect the expression of the LBcpLB tension probe or Lamin B1 (sFig. 2D) or the inherent structure of cytoskeletons (Fig. 4C and 4J). Under hypotonic conditions, IF tension increased when myosin and kinesin were inhibited and decreased when dynein was inhibited (Fig. 4D, 4E, 4F and 4G). Meanwhile, IF tension also elevated when the two motors of MT were inhibited (Fig. 4F and 4G). The data suggests that IF tension is modified synergistically by inward force of MF and MT in response to hypotonic stimuli.

**Figure 4.**
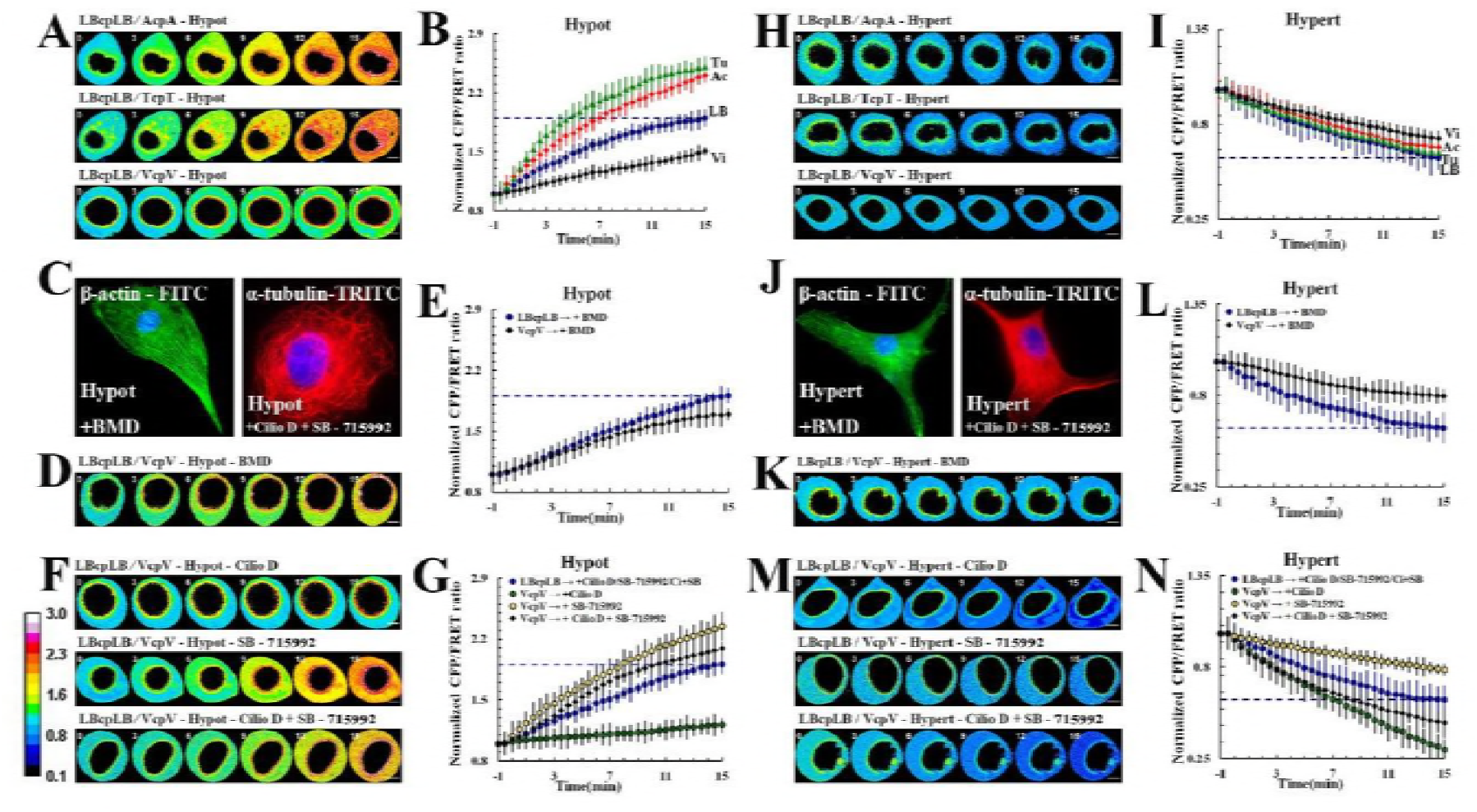
cpstFRET measurements on the influence of molecular motor inhibitors on cytoskeletal structural tension and nuclear lamina tension in response to osmotic pressure. The cpstFRET probe module was used to detect changes in cytoskeletal and nuclear lamina tensions in MCF-7 cells. Representative images of normalized CFP/FRET ratios subjected to myosin (BMD), dynein (Cilio D) or kinesin (SB-715992) inhibitors under hypo- or hyper-osmotic pressure. Under hypotonic conditions: (A) NL, MF, MT, and IF tensions were analyzed in MCF-7 cells (n = 8). (B) Normalized CFP/FRET signals corresponding to NL, MF, MT, and IF tensions versus time. (C) Left panel: MCF-7 cells were treated with BMD. β-actin were stained with FITC and the cells were also stained with DAPI to localize the nuclei. Right panel: MCF-7 cells were treated with Cilio D and SB-715992. α-tubulin were stained with TRITC and the cells were also stained with DAPI to localize the nuclei. Images were captured through confocal laser microscopy after immunofluorescence staining (n = 10). (D) NL and IF tension were analyzed in MCF-7 cells subjected to BMD (n = 9). Normalized CFP/FRET signals (E) compared to NL and IF tensions in Fig. 4D. (F) NL and IF tension were analyzed in MCF-7 cells subjected to Cilio D or SB-715992, respectively or simultaneously (n = 9). Normalized CFP/FRET signals (G), compared to NL and IF tensions in Fig. 4F. Under hypertonic conditions, (H) NL, MF, MT, and IF tensions were analyzed in MCF-7 cells (n = 9). (I) Normalized CFP/FRET signals corresponding to NL, MF, MT, and IF tensions versus time. (J) Left panel: MCF-7 cells were treated with BMD. β-actin were stained with FITC and the cells were also stained with DAPI to localize the nuclei. Right panel: MCF-7 cells were treated with Cilio D and SB-715992. α-tubulin were stained with TRITC and the cells were also stained with DAPI to localize the nuclei (n = 10). (K) NL and IF tension were analyzed in MCF-7 cells exposed to BMD. Normalized CFP/FRET signals (L) compared with NL and IF tensions in Fig. 4K (n=9). (M) NL and IF tension were analyzed in MCF-7 cells exposed to Cilio D or SB-715992, respectively or simultaneously. Normalized CFP/FRET signals (N) compared with NL and IF tensions in Fig. 4M (n=9). The calibration bar was set from 0.1 to 3.0. Scale bar, 10 μm. All error bars represent the SEM.

To follow, we challenged cells with hyper-osmotic conditions. As shown in figure 4H, MT-, MF-, IF-, and NL-tensions decreased, with the lowest NL tension in magnitude (Fig. 4H and 4I). Then, we suppressed the molecular motors. The IF tension was showing a trend of slight increase when BMD and SB-715992 were applied, whereas drooped obviously in response to Cilio D (Fig. 4K, 4L, 4M and 4N). Down-regulating of the MT tension by inhibition of dynein and kinesin simultaneously reduced IF tension dramatically (Fig. 4M and 4N). On the premise of making comprehensive consideration for the inward traction made by myosin and kinesin and the outward pulling driven by dynein, these data suggested that the MF and MT forces acted antagonistically to mediate IF tension by resisting the extracellular hypertonic.

In abnormal energy expenditure cells caused by DZ (calcium channel blockers) and NaN_3_ (mitochondria respiratory chain inhibitor), the tension of MFs and MTs were negligibly small, suggesting that their cytoskeletal structures alone do not generate tension (sFig. 6A and 6C). To summarize, our data suggests that the cytoskeleton could compete with inward or outward stresses caused by changes in corresponding ambient osmotic pressure. The three types of intracellular tension activities mediate IF tension differently. IF tension is adjusted synergistically or antagonistically by MF- and MT tensions against osmotic pressure.

Under hypotonic conditions, the normalized CFP/FRET ratio of NL tension increased to 1.9 levels. Intriguingly, irrespective of which molecular motor was inhibited, the normalized CFP/FRET ratio of NL tension reached to 1.9 invariably, in accordance with the control conditions (Fig. 4B, 4E, and 4G). We also observed the phenomenon that the normalized CFP/FRET ratio of NL tension decreased to 0.6 levels under hypertonic conditions, whereas the value remained unchanged in response to disparate molecular motor inhibitors (Fig. 4I, 4L, and 4N). The same phenomenon was observed in resting cells. Under isotonic conditions, the NL tension remained at a normalized CFP/FRET ratio of 1 in response to individual inhibitors, whereas the MF and MT tension altered correspondingly (sFig. 8A, 8B, 8C, and 8D). The results suggest that the NL tension associates little with MF and MT forces when respond to the changes of extracellular osmotic pressure, but influence by the outward or inward seepage pressure gradient.

### 4. The oxidative-induced increase of NL tension results from intracellular hyper-osmosis, associated closely with mass protein-nanoparticles production

Except for tension stimuli, chemical signals can also trigger morphological changes in the nucleus through intracellular tension activity. To gain better insight, we applied Adriamycin (ADM), H_2_O_2,_ and cycloheximide (CHX) to induce cellular senescence, oxidative stress, and apoptosis, respectively. These reagents had no impact on the amount of the LBcpLB tension probe (sFig. 2E). In terms of H_2_O_2_ application, both IF- and NL- tension increased markedly, while MF- and MT- tension remained unchanged (Fig. 5E and 5F). When Nesprin 3, a protein which links IFs with nuclear lamin, was disturbed, the NL tension notably declined compared with scrambled siRNA (Fig. 5G and 5H). Immunofluorescence assays showed that both MFs and MTs depolymerized in reaction to the H_2_O_2_ stimulus (Fig. 6A). Nanometre scale particle size observation and osmometry revealed the presence of nanometre-sized particles (Fig. 6B, 5–6 nm and 7–8 nm) and changes in intracellular osmotic pressure (Fig. 6C). Meanwhile, western blot analysis indicated that p-cofilin and p-stathmin levels significantly declined. The total actin and tubulin concentrations were unchanged (Fig. 6D). The data also suggest that H_2_O_2_ induces MF and MT depolymerisation and the production of mass protein-nanoparticles through cofilin and stathmin activation. Changes in IF tension results from intracellular osmotic pressure, particularly the colloid osmotic pressure, and NL tension was mediated primarily by IF tension under H_2_O_2_-induced oxidative stress cells.

**Figure 5.**
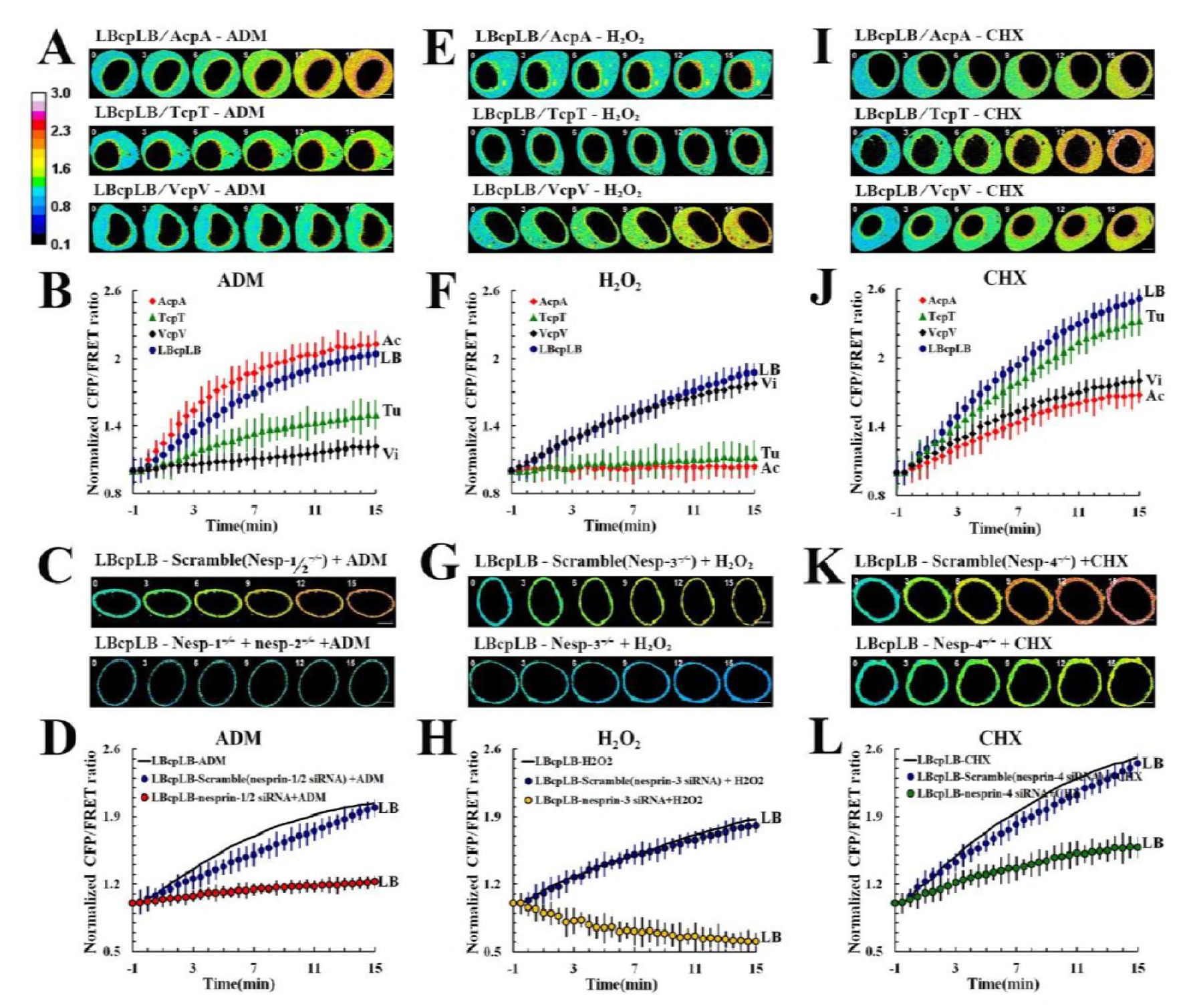
cpstFRET measured changes in cytoskeletal structural and nuclear lamina tensions in response to chemical stimuli. The NL, MF, MT, and IF tensions were analyzed in the senescence model of MCF-7 cells exposed to ADM alone (A, n = 8), and the NL tension was analyzed in response to ADM with Nesprin 1 and 2 (C, n = 8). (B) Normalized CFP/FRET signals corresponding to NL, MF, MT, and IF tensions versus time in response to ADM stimulation. (D) Normalized CFP/FRET signals corresponding to NL tension versus time in response to ADM stimulation with BMD (blue) and Nesprin 1 and 2 (red). The NL, MF, MT, and IF tensions were analyzed in the oxidative stress model of MCF-7 cells subjected to H_2_O_2_ exposure alone (E, n = 9), and the NL tension was analyzed alones in response to H_2_O_2_ with Nesprin 3 (G, n = 9). (F) Normalized CFP/FRET signals corresponding to NL, MF, MT, and IF tensions versus time in the presence of H_2_O_2_. (H) Normalized CFP/FRET signals corresponding to NL tension versus time in the presence of H2O2 with BMD, Cilio D, and SB-715992 (blue) and with Nesprin 3 (yellow). The NL, MF, MT, and IF tensions were analyzed in the apoptosis model of MCF-7 cells exposed to CHX alone (I, n = 9), and the NL tension was analyzed separately in the presence of CHX with Nesprin 4 (K, n=9). (J) Normalized CFP/FRET signals corresponding to NL, MF, MT, and IF tensions versus time in the presence of CHX. (L) Normalized CFP/FRET signals corresponding to NL tension versus time in the presence of CHX with Cilio D and SB-715992 (blue) and with Nesprin 4 (green). The calibration bar was set from 0.1 to 3.0. Scale bar, 10 μm. All error bars represent the SEM.

**Figure 6.**
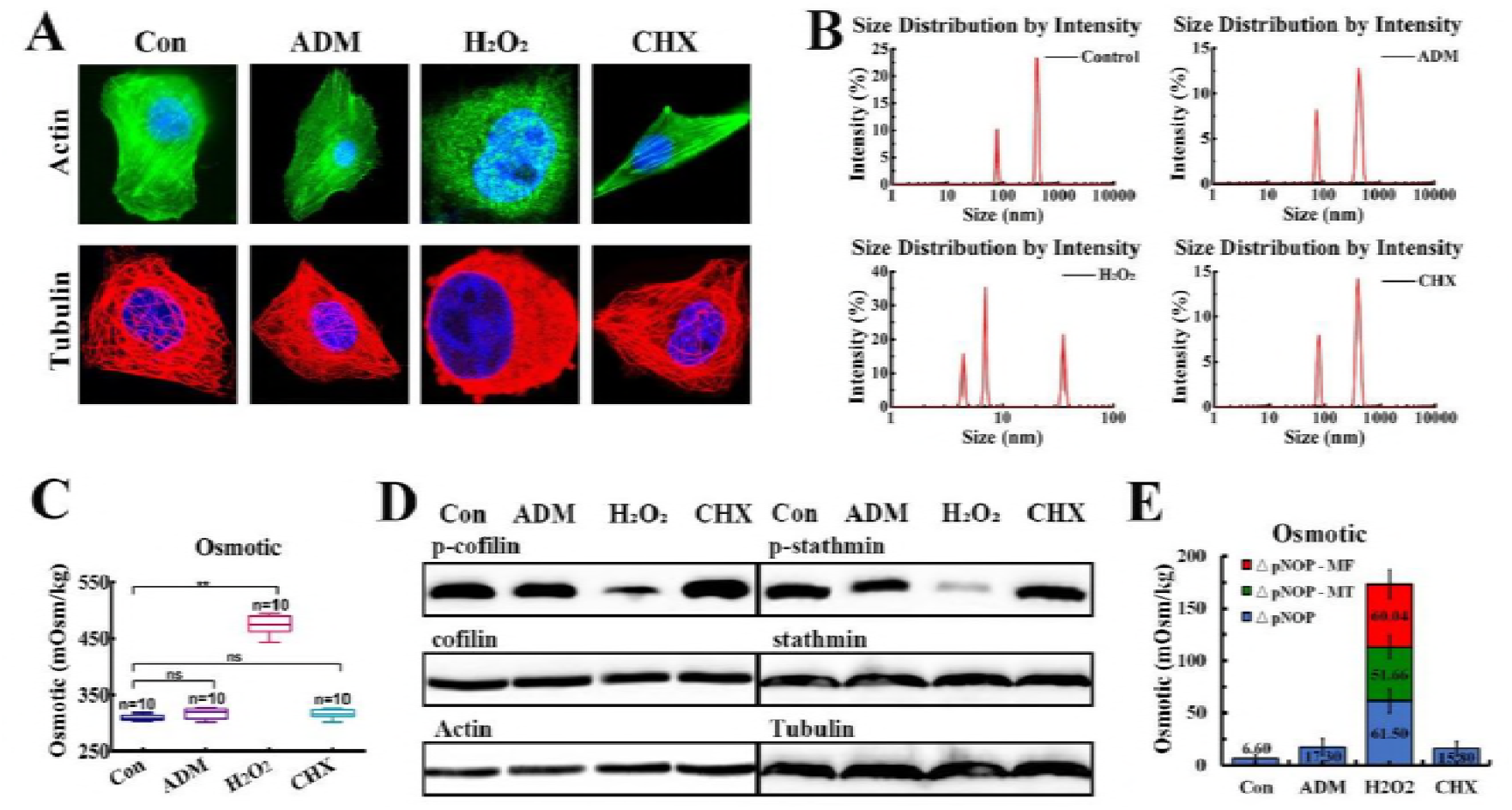
Intracellular osmotic pressure changes in response to various chemical stimuli. (A) MCF-7 cells were treated with ADM, H_2_O_2_, and CHX separately. The samples were stained for both β-actin and α-tubulin. The cells were also stained with DAPI to localize the nuclei. The images were captured through confocal laser microscopy after immunofluorescence staining (n = 10). (B) Intracellular nanoparticle size distribution in MCF-7 cells separately treated with ADM, H_2_O_2_, and CHX (n = 10). (C) MCF-7 cell intracellular osmotic pressure was measured via osmometery after the cells were exposed to ADM, H_2_O_2_, and CHX. (**: 0.001 < p < 0.05, ns: p > 0.05, Tukey-b test, n = 10). (D) P-cofilin, cofilin, actin, p-stathmin, stathmin, and tubulin levels in MCF-7 cells that had been treated with ADM, H_2_O_2_, or CHX (n = 6). (E) In the H_2_O_2_ group, the ΔpNOP could be divided into two parts by comparing the total intensity of the protein nanoparticles, generated from MF and MT depolymerization, respectively (versus Fig. 6B, line 1, row 2). Scale bar, 10 μm. All error bars represent SEM.

Then, we tested cells after the application of ADM. NL and MF tensions increased significantly while MT and IF tension only slightly increased (Fig. 5A and 5B). When we suppressed Nesprin 1 and 2, the proteins that link MFs with nuclear lamina, were disturbed, the NL tension declined from the original value compared with scrambled siRNA transfected cells (Fig. 5C and 5D). ADM induced no obvious changes in MF or MT structures (Fig. 6A). Furthermore, no protein-nanoparticles were produced obviously (Fig. 6B) and no apparent changes in p-cofilin and p-stathmin expression were observed in the presence of ADM (Fig. 6D).

After CHX treatment, NL and MT tensions markedly increased, while MF and IF tensions only slightly increased (Fig. 5I and 5J). Nesprin 4 disruption resulted in a further decline in NL tension compared with scrambled siRNA transfected cells (Fig. 5K and 5L). CHX stimulus resulted in non-significant changes in intracellular osmotic pressure (Fig. 6C). CHX had no obvious influence on MF or MT structures (Fig. 6A) and elicited no protein-nanoparticle generation (Fig. 6B). Moreover, CHX stimulation induced no obvious changes in the expression of p-cofilin and p-stathmin (Fig. 6D and 6E).

Although ADM and CHX stimuli non-significantly increased intracellular osmotic pressure, the balance of intracellular colloid-related and ionic osmotic pressure changed (Fig. 6E). Moreover, the NL tension had little concern with MF and MT tension when answer to the increased intracellular osmotic pressure caused by H_2_O_2_ treatment, further complementally illuminated and confirmed the role of NL in sensing changes of osmotic pressure.

## Discussion

Nucleus as the largest and stiffest organelle in the cell constitutes a subjectival structure to cell stabilization. Deformation of nucleus is dependent upon interactions between nuclear lamina tension and cytoskeleton tension, which is involved in a variety of cell activity, such as nuclear division, pyknosis, and migration. Fluorescence resonance energy transfer (FRET) – based tension sensors, dependent on the angular orientation of the donor and acceptor, have enabled the spatial and temporal analysis of mechanical forces across nuclear lamina and cytoskeleton in living cells. Because the angular sensor “hangs” on the cytoskeleton, it has less of an impact on the form and function of the intracellular skeleton structure (Fig. 1B and S. Table 1). Isolated nuclei respond to external gradient osmotic pressure, and NL tension can be evaluated quantificationally using the Laplace pressure formula (sFig.4). LBcpLB tension sensors have been found to be most sensitive from 1 pN to 12.59 pN. In the living cells, the NL counters a resting strain of the ~4.26pN. The existence of mechanical force on the nuclear lamina is supported by a number of previous studies that have observed nuclear deformation in response to artificial pull forces^43-45^. The study of cytoskeleton tension sensors suggests that variation in constitutive nuclear lamina tension originates from interactions of strains from MFs, MTs, and IFs. Remodeling of the MF, MT, or IF forces also drive changes in nuclear lamina tension and nucleus deformation.

To further untangle the strong interplay between the NL and cytoskeleton tensions, the present study requires the elimination of certain cytoskeletal structural tensions while other cytoplasmic mechanosensitive pathways keep intact. However, depolymerization of MFs or MTs can lead to the generation of protein nanoparticles. The damage to cytoskeletons is not merely the breakage of cytoskeleton structures and the elimination of structural forces, but is also the creation of the outward colloid osmotic pressure elicited by protein-nanoparticles. Therefore, the creation of the osmotic pressure contributes to enhancements in IF and NL tensions. Apart from intracellular force magnitudes, their vectors are crucial factors in subcellular tension activities. The hypo-osmotic pressure induces swelling pressure to plasma membrane, transmitting to NL through outward pull forces of IFs. Unlike IFs, the vectors of MF- and MT-dependent tensions are dependent upon the polarity of their structures. In response to hypo-osmotic pressure, direction of MF tension is inward due to its plus ends adjacent to the membrane and myosin moves to these plus ends. Similarly, direction of MT tension also is mainly inward because of kinesin is activated and walks to the plus ends adjacent to the membrane. However, dynein exerts different function on MT tension according to its move to minus ends and reverse force direction. So that alleviation of inward MT and MF forces promotes the increment of IF elicited by extracellular hypo-osmotic pressure.

In response to hyper-osmotic pressure, the compressive stress can reduce cytoskeleton tension and result in inward IF tension. IFs serve as “pistons” radiating from the nucleus and extending to the plasma membrane, and their tension activities are also dependent upon interactions with MF and MT forces. MF forces had little impact on IF tension, whereas outwards MT forces are involved in regulation of IF tension response to activation of dynein but not kinesin under an extracellular hypertonic circumstance. It can be speculated that hyper-osmotic stimuli induce shrinking pressure to plasma membrane, transmitting to NL through the inward compressive forces of IFs, and producing the weakened NL tension. Therefore, alteration of IF tension results from interactions among osmotic pressure, MF and MT forces, in which tension vector exerts an essential function.

Vector of intracellular tension activity can be regulated, involving crucially in cell polarity and nucleus deformation. It has been known widely that vector of MT force can be modified by the motor molecules dynein and kinesin due to their different walking directions. However, it is difficult for the MF force to change its vector, which is controlled strictly by myosin motivation and polar structure of MFs. Our studies found that the depolymerization of MFs or MTs and the generation of protein-nanoparticles led to a productive of outward oncotic pressure and swell of membrane, which served to compete with IF tension for nucleus deformation. Therefore, the production of protein-nanoparticles can exert essential function in the conversion of direction of intracellular tension, especially inward MF forces. Mechanism underlying vector of intracellular tension activity could provide a reasonable explain to cell polarity and deformation, like neuron polarity, astrocyte swelling, tumor cell invasion and metastasis, and so on^46^.

NL tension is a consequence of the resultant force of MFs, MTs, and IFs; however, it is unreasonable to decipher the alteration of NL tension according to the vector relationship between the cytoskeleton tensions mentioned above. Since vector of MF and MT forces depend on the polarity between the membrane and the cytoplasm, a logical model can be constructed, thus, MF or MT pulling forces can be produced simultaneously from the plasma membrane to the cytoplasm and from the nuclear membrane to the cytoplasm. The MFs near plasma membrane produce inward tension on IFs due to myosin walk to plasma membrane. But the MFs near nuclear envelope could lead to outward MF tension on NL because of myosin walk to nuclear envelope. Therefore, the inward and outward tension activity of MFs is counteracted through transmission of the IF structure, which attenuates IF tension and executes the transmission of extracellular osmotic pressure to the NL. So was the polarized structure of MTs and its tension activities (Table.1). The models suggesting that the transmission of osmotic pressures on NL tension occurs via IFs, associated closely with highly polarized MF and MT structures and their vectorially force activities.

**Table. 1.** Interaction of osmotic pressure with MF, MT, IF and NL-dependent tensions

| Treatment | Parameter | Osmotic pressure | membrane |  | IF tension | nucleus |  | NL tension | Phenomenon |
| --- | --- | --- | --- | --- | --- | --- | --- | --- | --- |
|  |  |  | MF | MT |  | MF | MT |  |  |
|  |  |  | tension | tension |  | tension | tension |  |  |
| Hyperosmotic pressure | Magnitude | ↑ | ↑ | ↑ | ↑ | ↑ | ↑ | ↑ | Swell |
|  | Vector | ← | → | → | ← | ← | ← | ← |  |
| Hyperosmotic pressure | Magnitude | ↑ | = | ↓ | ↓ | = | ↓ | ↓ | Shrink |
|  | Vector | → | → | ← | → | ← | → | → |  |

\*↑ indicates the increment; ↓ indicates the reduction;

The present study shows that NL tension can hold steady, be increased or decreased in responses to extracellular iso-, hypo- or hyper- osmosis pressure. However, changes in NL tension are not dependent on activity of MF and MT force, suggesting that NL tension can be controlled fatefully by cellular osmosis pressure though NL exists in center of a cell. Meanwhile, intracellular protein-nanoparticle-related osmotic pressure results from depolymerization of the MFs or MTs, and also results in enhanced NL tension. NL tension appears to be of the consistent order of magnitude in response to alteration of osmotic pressure. The data suggest that NL serves as the osmolarity sensor. The phenomenon can be explained reasonably by the models mentioned above. Alteration in the IF tension is dependent mostly on MF and MT strains, and a change in NL tension is linked to extra- or intra-cellular OP. Therefore, intracellular NL, but not IFs, cloud be treated as the osmolarity sensor, detecting extracellular and intracellular OP remotely and accurately.

NL tension in living cells can also be modified by chemical stimuli, involving in senescence, apoptosis and oxidative stress. Alternation in NL tension is associated closely with intracellular steering force activity. MF forces result from senescence inducers, MT force are elicited by apoptosis stimulators, and IF tension is induced by oxidizing agents, all of which can exert a tension effects effectively on NL. It should be point out that oxidant-induced IF tension results from upregulation of intracellular osmotic pressure, in which oxidant activates MF and MT depolymerize factors cofilin and stathmin and induces production of protein-nanoparticles and creates outward OP, performs vector conversation of intracellular tension activity. Therefore, the production of protein-nanoparticles may be associated closely with the occurrence and development of oxidative-related diseases.

The role of intracellular structural force on NL tension is dependent upon transmission of the LINC complex in response to chemical signals. MF, MT, and IF forces exert their individual function on NL through difference of LINC complex, involving in nesprin-1, -2, -3, and -4. Knocking down different types of nesprin (-1 and -2, -3, and -4) can disrupt the physical correspondence connections between MFs, MTs or IFs to NL, and cut off force transmissions of different cytoskeletons to NL. The LINC complex proteins function as the core mediator for the transmission of cytoskeletal forces to NL, which is different obviously from OP-induced NL tension and their underlying mechanisms (Table.1). The data suggest that intracellular force activity in NL is regulated by structural forces and adapt protein of cytoskeleton together in response to chemical signals. In addition, chemical stimuli can result in alteration of osmotic pressures, including its composition and magnitude. The data highlights the significance of distinguishing the composition of intracellular osmotic pressure, which will bring new insights into the cellular stress responses in physiological and pathological conditions.

The present study indicates that there are five steerable forces that control intracellular tension activities, namely ionic and protein-nanoparticle related osmotic pressures, MF forces, kinesin and dynein-dependent MT tension, which interact to control intracellular NL tension, both in terms of magnitude and vectors. Intracellular protein-nanoparticle related osmotic pressure acts as a vector converter and contributes to the outward expansion of NL via the passive pull of Ifs, and the depolymerization of MFs or MTs, and the production of mass protein-nanoparticles. Meanwhile, NL can remotely detect extra- and intra- cellular osmosis pressure alterations, which are closely associated with the highly polarized structures of MFs and MTs and their relative tension activities. Oxidative stress results in the increase of NL tension, associated closely with protein-nanoparticles-related osmotic pressure and cofilin and stathmin activation. Interactions between nucleoskeletal and cytoskeletal tensions depend on the transmission of cues from LINC complex proteins in response to chemical stimuli. The newly-established analytical system can explore the subcellular mechanical activities reasonably, including those of the nucleus, which improves and deepens the understanding of mechanisms underlying intracellular mechanical tension, and could therefore develop new insight into the elucidation of tension-related pathogenic mechanism, identification of new drug targets, and development of therapeutic drugs and strategies.

## Acknowledgements

We thank Frederick Sachs and Fanjie Meng (University at Buffalo) for providing the plasmids encoding actin-cpstFRET (AcpA) and actin-cpstFRET (Acp). This work was supported by grants from the National Natural Science Foundation of China (81573409), the Natural Science Foundation of Jiangsu Province (BK20161574), and the Project of Priority Academic Program Development of Jiangsu Higher Education Institutions (Integration of Traditional Chinese and Western Medicine).

## Author contributions

Jun Guo was involved in the initial conceptualization of the project. TingTing Chen and Huiwen Wu wrote the manuscript with input from all authors. TingTing Chen performed and analysed experiments. YuXuan Wang and JiaRui Zhang performed osmotic pressure and western blot test. JinJun Shan and Huanhuan Zhao helped with manuscript preparation. Jun Guo initiated and supervised the project.

## MATERIALS AND METHODS

### 1. Reagents and antibodies

Cytochalasin D (cyto D), nocodazole (Noc), Doxycycline (Doxy), NH4Cl and H_2_O_2_ were obtained from Aladdin (Shanghai, China). Adriamycin (ADM), cycloheximide (CHX), and diltiazem (DZ) were purchased from Sigma-Aldrich (St. Louis, Missouri, USA). Sodium azide (NaN_3_) and 2,3-Butanedione-2-monoxime (BMD) were acquired from Santa Cruz (CA, USA). Cilliobrevin D (Cilio D) was purchased from Merck Millipore (Darmstadt, Germany). Ispinesib (SB-715992) was procured from BioVision (California, USA). FITC-Phalloidin was obtained from Solarbio (CA1620, Shanghai, China). Antibodies were purchased from commercial sources: rabbit anti-lamin B1 antibody (13435S, Cell Signaling Technology, Temecula, USA), rabbit anti-β-actin antibody (4970P, Cell Signaling Technology, Temecula, USA), mouse anti-tubulin-α antibody (T5168, Boster, Wuhan, China), rabbit anti-vimentin antibody (5741S, Cell Signaling Technology, Temecula, USA), rabbit anti-GFP antibody (2956S, Cell Signaling Technology, Temecula, USA), mouse anti-phospho-cofilin (Ser24) (bs-10252R, Bioss, Woburn, USA), mouse anti-cofilin antibody (PB0035, Boster, Wuhan, China), rabbit anti-phospho-stathmin 1 (Ser38) (bs-3432R, Bioss, Woburn, USA), and mouse anti-stathmin 1 antibody (BA1417, Boster, Wuhan, China).

### 2. Cell Culture

MCF-7 (human breast metastatic adenocarcinoma) cells (HTB 22, ATCC, Manassas, USA) were cultured on fibronectin coated coverslips in Dulbecco’s Modified Eagle Medium (DMEM, 319-005-CL, Wisent, Montreal, Canada) containing 10% fetal bovine serum (086-150, Wisent, Montreal, Canada) and a 1% mixture of penicillin and streptomycin (Gibco, Invitrogen, USA) at 37°C in 95% air and 5% CO_2_. Cell morphology and growth were monitored and the cells were routinely tested for mycoplasma contamination using a Mycoplasma Plus PCR Primer Set (Agilent, Santa Clara, CA, US) and all cells used were found to be negative for mycoplasma.

### 3. Tension Sensor Design

The sensors were designed using the NovoRec^®^ PCR Seamless Cloning Kit and restriction enzyme cloning techniques, in accordance with previous reports^39, 48^. Based on Venus and Cerulean, we constructed circularly permutated cpVenus and cpCerulean. Seven amino acids were added between them to generate cpstFRET model (cpVenus-3Gly-Pro-3Gly-cpCerulean)^49, 50^. Then, we substituted Venus and Cerulean with cpVenus and cpCerulean, so that circularly permutated variants generated cpstFRET. The nuclear laminB1 is a V intermediate filament protein featuring a central α-helical rod domain that is flanked by non-helical head and tail domains. Then, we cloned lamin B1 and incorporated the EGFP gene at the C-terminal of cpstFRET, creating pCMV-laminB1-cpstFRET (Lcp). We then inserted another lamin B1 gene into the C-terminal of cpstFRET and created pCMV–laminB1-cpstFRET-laminB1(LcpL). Additionally, we designed pCMV-actin-cpstFRET (Acp), pCMV-actin-cpstFRET-actin (AcpA), pCMV-tubulinα-cpstFRET (Tcp), pCMV-tubulinα-cpstFRET-tubulinβ (TcpT), pCMV-vimentin-cpstFRET (Vcp) and pCMV-vimentin-cpstFRET-vimentin (VcpV) sensors.

### 4. Plasmid Extraction and Transfection

The E.Z.N.A Endo-free plasmid DNA Mini Kit II (OMEGA, Tarzana, USA) was used to extract single colony plasmids, according to the manufacturer’s instructions, after which, the plasmids were stored at −20 °C. The cells were then transfected for 24 h before the experiments with tension sensor plasmids using FuGENE 6 Transfection Reagent (Roche Diagnostics, Basel, Switzerland), following the manufacturer’s specifications. The transfection efficiency was approximately 82.5%. The transfected MCF-7 cells were sorted using a Moflo XDP cell sorter with Summit 5.3 software (Beckma Coulter). Samples were analyzed using the following excitation lasers and emission filters: Cyan-458-nm laser and 447/60 bandpass filter; Yellow-514-nm laser and 580/23 bandpass filter. Cells were sorted when the cyan and yellow emission wavelengths were detected at the same time. The sorted MCF-7 cells were diluted into single cells and then seeded on 96-well plates. In all experiments, the transfected and sorted single cell lines were trypsinized, the cells were then incubated in cell culture media for 24–36 h. Attached cells were gently washed by PBS before imaging.

### 5. cpstFRET analysis

The efficiency of FRET depends on the distance and the dipole angular orientation between the donor/CFP and the acceptor/YFP. Cells were imaged on a Leica confocal microscope SP5 equipped with a × 63 oil-immersion objective. The donor and acceptor were observed under 458 nm and 514 nm argon lasers, respectively. CFP/FRET ratios were calculated using the equation 1/R = Cerulean donor/Venus acceptor.

### 6. FRET-AB and FRAP analysis

The donor excitation light was set to 10%, the acceptor detection channel was opened and adjusted to the appropriate excitation light intensity. The ROI (region of interest) was then selected and the acceptor components within the cells were bleached, and the efficiency of FRET was calculated. FRAP was used to observe the diffusion of molecules and bleaching of fluorescently labeled proteins inside cells with a high intensity laser pulse. When proteins were transiently bound to structures in the photo bleached area of laminB1, the fluorescence recovered through exchange between the fluorescently labeled diffused molecules and the bound photo bleached molecules. The recovery curve can be used to estimate the protein flow rate.

### 7. Nuclear extraction in vitro

Cells (transfected with tension sensors) were placed in an ice-bath and the cells were washed with cold PBS. I cold Nuclei EZ lysis buffer (SIGMA-ALDRICH) was then added to each dish and the cells were scraped into centrifuge tubes, according to the manufacturer’s instructions. The isolated nuclei were balanced and cultured (5 × 10^4^ cells/cm^2^) in complete media for 2 h prior to the next treatment.

### 8. Model of isolated nuclear as the thin-film according to the theory of membrane

Buffer movement through the nuclear lamina caused nuclear envelope swelling or shrinking. According to the non-moment theory of shells, the nuclear lamina tension components subjected to pressure (internal or external) were:

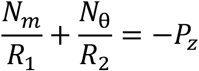

In the horizontal section, the changes in radial force created by nuclear laminar surface tensions was equivalent to permeable pressure between the inner nucleus and the outside:

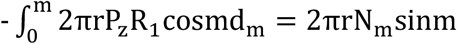

To assess the various pressure microelements on the nuclear lamina, we calculated the nuclear lamina tension via Laplace pressure calculations. The microelemental pressure on the nuclear lamina was comprised of both circumferential and axial stresses. The circumferential stress was much greater than the axial stress. The main horizontal compressive tangent maintained nuclear laminar tension, however, the vertical tangent stress can be ignored as it was insignificant. In this calculation, the Law of Laplace relates membrane tension to trans-membrane pressure and the radius of the nucleus radius.

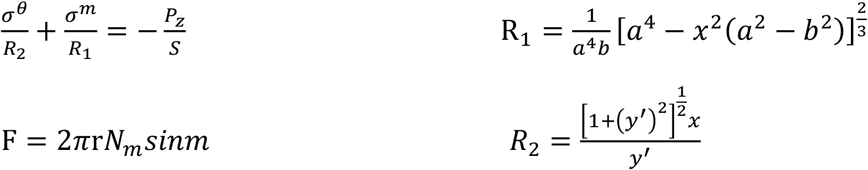

Uniform distribution of permeable pressure differences resulted in equal effective stress on the nuclear lamina elements.

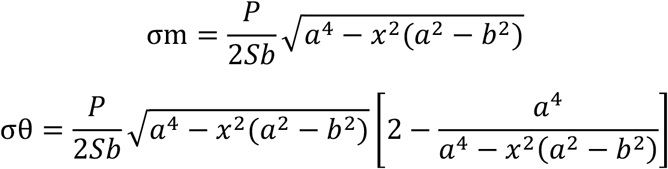

The formula for the tension calibration curve in nuclear lamina is:

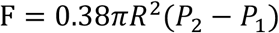

### 9. FLIM analysis

Time-domain fluorescence lifetime imaging (FLIM) was performed using the time-correlated single-photon counting technique (TCSPC). FLIM experiments and FLIM data analysis were performed as described previously using FLIM acquisition software (SPCImage, Becker & Hickl). Fluorescence lifetime was determined by fitting a single-exponential decay model, as LBcpLB modification has two pixels. ECFP was excited with a mode-locked laser tuned to 405 nm and eYFP was excited with a mode-locked laser tuned to 515 nm. Photons were counted and correlated with excitation laser pulses using a Becker and Hickl DCS-120 module (Berlin, Germany). Acquisition times of up to 500 s achieved sufficient photon statistics.

### 10. Osmotic pressure evaluation

Osmotic pressure exerts outward tensile force on the cytoskeleton and changes the morphology of cells. We used 16 different salt solution concentrations to create different osmotic pressure environments for MCF-7 cells (varying between 0 mOsm kg^−1^ and 600 mOsm kg^−1^), which were measured using a freezing point osmometer (Osmomat 030, Gonotec).

### 11. siRNA transfection

MCF7 cells were transfected with siRNA targeting nesprin-1, nesprin-2, nesprin-3, or nesprin-4 genes (Sangon Biotech, Shanghai, China) using FuGENE 6 Transfection Reagent (Roche Diagnostics, Basel, Switzerland).

### 12. Western Blot

Proteins were extracted from cells using SDS/PAGE. Then, the protein band was transferred to the NC membrane via wet electrotransfer. The main antibodies and dilutions used were: rabbit anti-lamin B1 antibody (CST; 1:1000), rabbit anti-β-actin antibody (CST; 1:1000), mouse anti-tubulin-α antibody (T5168, Boster; 1:300), rabbit anti-vimentin antibody (CST; 1:1000), rabbit anti-GFP antibody (CST; 1:1000), mouse anti-phospho-cofilin (Ser24) (Bioss; 1:300), mouse anti-cofilin antibody (T5168, Boster; 1:300), rabbit anti-phospho-stathmin 1 (Ser38) (Bioss; 1:300), and mouse anti-stathmin 1 antibody (T5168, Boster; 1:300). After blotting, the protein side of the membrane was exposed to an X-ray film.

### 13. Immunofluorescence

In the immunocytochemical studies, MCF7 cells were washed twice and fixed in freshly prepared 4% paraformaldehyde solution for 20 min. The membrane was permeabilized in 0.1% Triton X-100 at 4 °C and blocked with 3% BSA/PBS for 30 min. For fluorescence labeling, cells were incubated with the primary antibody. After incubation overnight at 4 °C, the cells were washed and incubated with the secondary antibodies in darkness. DAPI was used to stain the nuclei. Labeled samples were examined by immunofluorescence using a confocal microscope (TCS SP5, Leica).

### 14. Analysis of particle-size distributions and colloid osmotic pressures

Nanoparticle-size distribution and agglomeration within supplemented cell medium was measured by dynamic light scattering (DLS) using a Zetasizer nano ZS90 (Malvern, UK). The count rate was measured through an angle of 0^0^ DLS using a Zetasizer nano ZS90 (Malvern, UK).

The molecular weight (M_W_) and the second viral coefficient (A_2_) of the supernatant were evaluated using a Zetasizer NanoS system that provides a method for measuring the molecular weight of the proteins at only one angle (0^0^) by using the Rayleigh equation – i.e. the relationship between the intensity of the scattered light of macromolecules and their weight-average molecular weight (M_W_). A plot of KC/R_θ_ versus C is expected to be linear with an intercept equivalent to 1/M and a slope equal to the second viral coefficient A2. The following relations (Rayleigh equation) are valid:

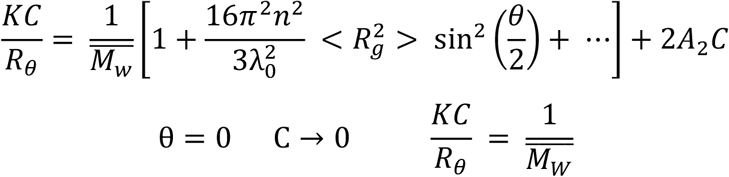

where: K – optical constant, C – polymer concentration, R_θ_ – Rayleigh ratio of the sample, M_W_ – molecular weight, A_2_ – 2^nd^ viral coefficient, θ – measurement angle. We appeal to the elegant statistical thermodynamic apparatus of our scientific forefathers, which relates macroscopic quantities (osmotic pressure) to molecular properties (molar mass). With the usual notation, the following relations (Van’t Hoff equation) are valid:

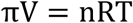

Deviations from ideality, normally studied in physical chemistry but at least mentioned in a beginning course, can be handled with a viral expansion:

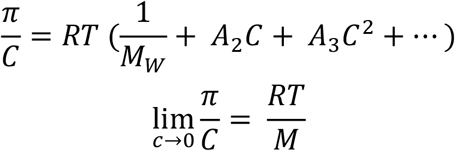

A plot of π/C versus C is expected to be linear with an intercept equivalent to M, thus the count rate is expected to be linear with osmotic pressure.

Thus, the count rate is linearly related to the osmotic modulus:

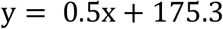

where: y – optical constant, x – count rate

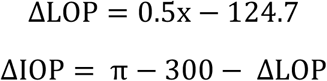

### 15. Data analysis

The images from CFP and YFP channels were aligned and the ratio of YFP/CFP was calculated using Image-J software. The FRET ratio in each subcellular region was measured for each cell and averaged over multiple cells. All data are expressed as Means ± SD or ± SEM. One-way ANOVA for single factor sample comparisons and LSD for comparisons between any two means. Each experiment was repeated at least three times, and > 10 cells were imaged and each condition was analyzed. The R of MCF-7 is presented as pixel count distribution from > 10 MCF-7 cells.

**Supplementary Table 1.** Percentage of apoptotic cells and cell cycle analysis for MCF-7 cells transfected with different probes. Cell apoptosis in MCF-7 cells transfected with LBcpLB, LBcp, AcpA, Acp, TcpT, Tcp, VcpV or Vcp plasmids. These data indicate that most cells transfected with the plasmids were not apoptotic, similar to controls. Cell cycle analysis, remaining in the G1/S phase. Additionally, the percentage of cells in the G2 and G1/G2 phases were low. Therefore, the plasmids had minimal influence on apoptosis and cell cycle progression. Stable cell lines expressing these probes showed unaffected cell physiology (n = 10; tissue culture plates were used as controls).

|  | Apoptosis | Cell cycle |  |  |  |
| --- | --- | --- | --- | --- | --- |
|  |  | G1 | S | G2 | G1/G2 |
| Con | 0.11 ± 0.05% | 41.43 ± 1.17% | 50.66 ± 1.11% | 7.91 ± 0.16% | 1.87 |
| LBcpLB | 0.08 ± 0.03% | 43.57 ± 1.37% | 51.60 ± 1.61% | 4.83 ± 2.76% | 1.89 |
| LBcp | 0.15 ± 0.06% | 42.31 ± 2.37% | 52.04 ± 2.12% | 5.65 ± 1.89% | 2.00 |
| AcpA | 0.19 ± 0.03% | 46.86 ± 2.07% | 47.14 ± 2.09% | 6.00 ± 2.48% | 1.92 |
| Acp | 0.05 ± 0.02% | 41.13 ± 1.97% | 50.56 ± 2.11% | 8.31 ± 1.86% | 1.91 |
| TcpT | 0.09 ± 0.04% | 42.64 ± 0.85% | 52.69 ± 2.13% | 4.67 ± 2.84% | 2.00 |
| Tcp | 0.12 ± 0.08% | 43.35 ± 2.06% | 52.28 ± 1.82% | 4.37 ± 1.77% | 1.91 |
| VcpV | 0.18 ± 0.06% | 45.56 ± 1.89% | 50.34 ± 2.14% | 4.10 ± 1.67% | 1.79 |
| Vcp | 0.16 ± 0.04% | 44.76 ± 2.55% | 48.13 ± 2.31% | 7.11 ± 2.49% | 2.02 |

**Supplementary Figure 1 | Localization of LBcpLB, AcpA, TcpT, and VcpV probes constructs.** Lamin B1, β-actin, α-tubulin and vimentin were individually stained. The cells were also stained with DAPI to localize the nuclei. MCF-7 cells were transfected with LBcpLB, AcpA, TcpT, and VcpV plasmids. Specific antibodies were employed to immunostain the MFs, MTs, IFs, and NLs respectively using antibodies against β-actin, α-tubulin, vimentin, and lamin B1. The images were generated via confocal laser microscopy after immunofluorescence staining (n = 10). Scale bar, 10 μm.

**Supplementary Figure 2 | Verification of tension probes and biochemical analysis of chemical agent stimuli on tension probes.** (A) Western blot of MCF-7 cells transfected with LBcp, AcpA, LBcpLB, VcpV or TcpT plasmids all expressing GFP mutant. (B) Western blot of MCF-7 cells transfected with LBcp, AcpA, LBcpLB, VcpV or TcpT plasmids expressing α-tubulin (row 1), β-actin (row 2), lamin B1 (row 3) or vimentin (row 4). (C) GFP mutant in transfected LBcp cells after the addition of MF and MT depolymerization agents (Noc and Cyto D, respectively) were compared with LBcpLB cells. (D) Expression levels of GFP mutant and Lamin B1 in transiently transfected LBcpLB cells treated with BMD, Cilio D, and SB-715992. (E) GFP mutant in transfected LBcpLB cells upon exposure to ADM, H_2_O_2_, and CHX respectively, compared to LBcp cells. (F) Expression levels of GFP mutant and Lamin B1 in transiently transfected LBcpLB cells treated with Nesprin 1, 2, 3, and 4 simultaneously or separately (n = 6).

**Supplementary Figure 3 | LBcp, Acp, Tcp, and Vcp negative control cells in response to different osmotic pressure states.** Representative 3D reconstruction of LBcp from bottom to top (A, line 1, row 1), Acp (B, line1, row 1), Tcp (C, line1, row 1), and Vcp (D, line1, row 1) cell (n = 12). Three-dimensional reconstruction spatial stereogram of negative control probes versus sFig. 3A-D line 1, row 1, respectively. Normalized CFP/FRET signals from LBcp, Acp, Tcp, and Vcp in hypotonic and hypertonic conditions (n = 8) versus time, versus sFig. 3A-D line 1, row 2 and 3, respectively. The calibration bar was set from 0.1 to 3.0. Scale bar, 10 μm. All error bars represent the SEM.

**Supplementary Figure 4 | FRET efficiency analysis of tension probes and the quantification of LBcpLB probes through isolated nucleus model experiments.** Experimental set-up for single-nuclear-lamina force-FRET measurements. The nucleus was extracted in vitro, and then the quiescent conditions of the nuclear lamina were observed (mean = 1). The nucleus suffered from osmotic pressure differences across the nucleus under these conditions. (A) The reliability of LBcpLB, AcpA, TcpT, and VcpV probes was tested by FRET acceptor photobleaching (n = 10). (B) and (C) Isolated nuclei that were stored in nuclei PURE storage buffer (600 mOsm kg^−1^) took 150 min to achieve the simulation conditions of normal cells in 300 mOsm kg^−1^ hepers solution (n = 12). CFP/FRET LBcpLB signals in response to hepers solution versus time. (D) and (E) After simulated conditions were achieved, the isolated nuclei were exposed to hypotonic conditions (differential: 300 to 0 mOsm kg^−1^; n = 12). LBcpLB CFP/FRET signal in response to isotonic conditions versus time. (F) After simulated conditions were achieve, the nuclei were treated with 16 different salt concentrations generated different osmotic pressures outside and inside the membrane (n = 12). The 16 different osmotic pressure differentials varied from between 300 mOsm kg^−1^ and 0 mOsm kg^−1^. Scale bar, 10 μm. The calibration bar was set from 0.1 to 3.0. All error bars represent the SEM.

**Supplementary Figure 5 | Model of isolated nuclei construction as the thin-film according to the theory of membrane.** (A) Representative 3D reconstruction of LBcpLB (FRET images) from bottom to top (n = 10). (B) Three-dimensional reconstruction spatial stereogram of LBcpLB probes versus sFig. 5A. In solution, the isolated nuclei were subjected to osmotic pressure only. Balance when the pressure difference was equal to the osmotic pressure. The nuclear lamina tension was equivalent to the permeable pressure. (C) Micro-elemental pressure on the nuclear lamina is comprised of both circumferential and axial stresses. Circumferential stress was much greater than axial stress. The vertical tangent stress may be ignored as it is too small to be of consequence against osmotic pressure. In consequence the main horizontal compressive tangent maintained the nuclear laminar tension. (D) Uniform distribution of permeable pressure differences resulted in equal effective stress on the nuclear lamina. Based on the analysis above, the tension calibration curve for LBcpLB is: F = 0.38πR^2^ (P_2_ - P_1_).

**Supplementary Figure 6 | Structural tension in MFs, MTs, IFs, and NLs in response to chemical stimuli.** The scatter diagrams show cytoskeleton and nuclear lamina under different chemical stimuli, containing (A) calcium channel blockers (DZ), (B) hypoxia (NH_4_CL), and (C) mitochondria respiratory chain inhibitor (NaN_3_; n = 8). Normalized CFP/FRET signal corresponding to structural tension versus time. The calibration bar was set from 0.1 to 3.0. Scale bar, 10 μm. All error bars represent SEM.

**Supplementary Figure 7 | Nanoparticle generation caused by MF and MT depolymerization.**

(A) MCF-7 cells transfected with LBcpLB were observed along the longitudinal section in response to exposure to Cyto D and Noc, respectively and simultaneously (n = 9). (B) MCF-7 cell intracellular nanoparticle dimensions and distribution after Cyto D and Noc treatment, respectively and simultaneously (n = 10). (C) Actin and tubulin levels in MCF-7 cells treated with Cyto D and Noc (n = 6).

**Supplementary Figure 8 | MF, MT, and NL structural tension alterations in response to molecular motor inhibitors.** The histograms depict microfilament, microtubule and nuclear lamina tensions upon the incorporation of three molecular motor inhibitors, containing (B) BMD, (C) Cilio D and (D) SB-715992 (n = 8). Normalized CFP/FRET signals corresponding to MF-, MT-, and NL-dependent tensions versus time. The calibration bar was set from 0.1 to 3.0. Scale bar, 10 μm. All error bars represent the SEM.

**Movie S1. Anisotropy ratio R of VcpV and LBcpLB-expressing MCF-7 cells subjected to treatment with Cyto D and Noc, simultaneously.** The video is presented using a 32-colour map (ImageJ software) with the same range as in Fig. 2C (line 1). Inset indicates the timing of drug administration.

**Movie S2. Anisotropy ratio R of AcpA and LBcpLB-expressing MCF-7 cells subjected to treatment with Cyto D and Noc, simultaneously.** The video is presented using a 32-colour map (ImageJ software) with the same range as in Fig. 2C (line 2). Inset indicates the timing of drug administration.

**Movie S3. Anisotropy ratio R of TcpT and LBcpLB-expressing MCF-7 cells subjected to treatment with Cyto D and Noc, simultaneously.** The video is presented using a 32-colour map (ImageJ software) with the same range as in Fig. 2C (line 3). Inset indicates the timing of drug administration.

